# High resolution ultrasonic neural modulation observed via *in vivo* two-photon calcium imaging

**DOI:** 10.1101/2021.03.23.436645

**Authors:** Zongyue Cheng, Chenmao Wang, Bowen Wei, Wenbiao Gan, Qifa Zhou, Meng Cui

## Abstract

Neural modulation plays a major role in delineating the circuit mechanisms and serves as the cornerstone of neural interface technologies. Among the various modulation mechanisms, ultrasound enables noninvasive label-free deep access to mammalian brain tissue. To date, most if not all ultrasonic neural modulation implementations are based on ∼1 MHz carrier frequency. The long acoustic wavelength results in a spatially coarse modulation zone, often spanning over multiple function regions. The modulation of one brain region is inevitably linked with the modulation of its neighboring regions. To significantly increase the spatial resolution, we explored the application of high-frequency ultrasound. To investigate the neuronal response at cellular resolutions, we developed a dual-modality system combining *in vivo* two-photon calcium imaging and focused ultrasound modulation. The studies show that the ∼30 MHz ultrasound can suppress the neuronal activity in awake mice at 100-micron scale spatial resolutions, paving the way for high-resolution ultrasonic neural modulation.

## Introduction

Neural modulation technologies hold great significance in neuroscience research and medical applications^1–6^. The capability of exciting or suppressing neuronal activity during behavior can reveal the functions of neural systems and enable brain-machine interfaces^5,7,8^. Among the various modulation mechanisms, ultrasound-based technologies allow a noninvasive label-free deep penetration in the mammalian brains, thanks to the weak attenuation of acoustic waves in brain tissue^9–17^. In the majority of applications, low frequency (∼ 1 MHz) ultrasound was employed^18–21^. Despite its wide adoption in neuroscience applications, a major drawback of the low-frequency ultrasound is its poor spatial resolution and confinement^20,22,23^. The near centimeter-scale modulation zone often covers multiple brain regions, far too coarse for neuroscience research^24^. For neural interface technologies, the large modulation zone also significantly limits the achievable degrees of control.

To drastically improve the spatial resolution and confinement, a straightforward solution would be to reduce the acoustic wavelength^25–28^. Thus, it would be of great significance to access the effect of high-frequency ultrasound on neurons and to investigate the frequency, magnitude, waveform required to achieve reliable neural excitation or suppression^29–38^. In this work, we explored the application of focused ultrasound with near 30 MHz carrier frequency. Under the same focusing condition, the 30 times higher frequency improves the spatial resolution by a factor of 30 in each dimension and reduces the 3D modulation volume by a factor of 27,000. To evaluate the neuronal activity within the greatly reduced modulation zone at high spatial resolutions, we developed a dual-modality system combining a polymer focused sound transducer and a two-photon fluorescence calcium imaging system, which allowed us to record the activity of neurons at cellular resolutions. Moreover, we employed EEG to monitor the response from neuronal populations.

## Results

The dual-modality system involves the combination of a two-photon laser scanning fluorescence microscope and a high-frequency focused ultrasound transducer (**Fig. 1a**). To accommodate the ultrasound transducer (PI35-2-R0.50, Olympus NDT), we employed a water-dipping objective lens (Nikon 16x NA 0.8) with a large access angle. We designed a water container that was attached to a large surface mouse head bar with an O-ring in between (**Fig. 1b**). The water immersion ensured a proper coupling to the brain tissue for both the light and the sound waves. To minimize the sound reflection, we utilized plastic coverslips as the optical cranial windows, which enabled ∼90% acoustic power transmission (**Supplementary Table 1**). Before *in vivo* imaging, we aligned the ultrasound transducer location such that the ultrasound focus was overlapped with the two-photon imaging focal plane. Experimentally, we mounted fluorescence beads in 2% agar as the calibration sample. The propagation of the ultrasound wave in agar led to a low-pressure zone which pulled the surrounding beads towards the ultrasound focus by hundreds of nanometers, allowing us to precisely visualize and align the ultrasound focus under the two-photon microscope (**Supplementary Fig. 1**).

**Figure 1.**
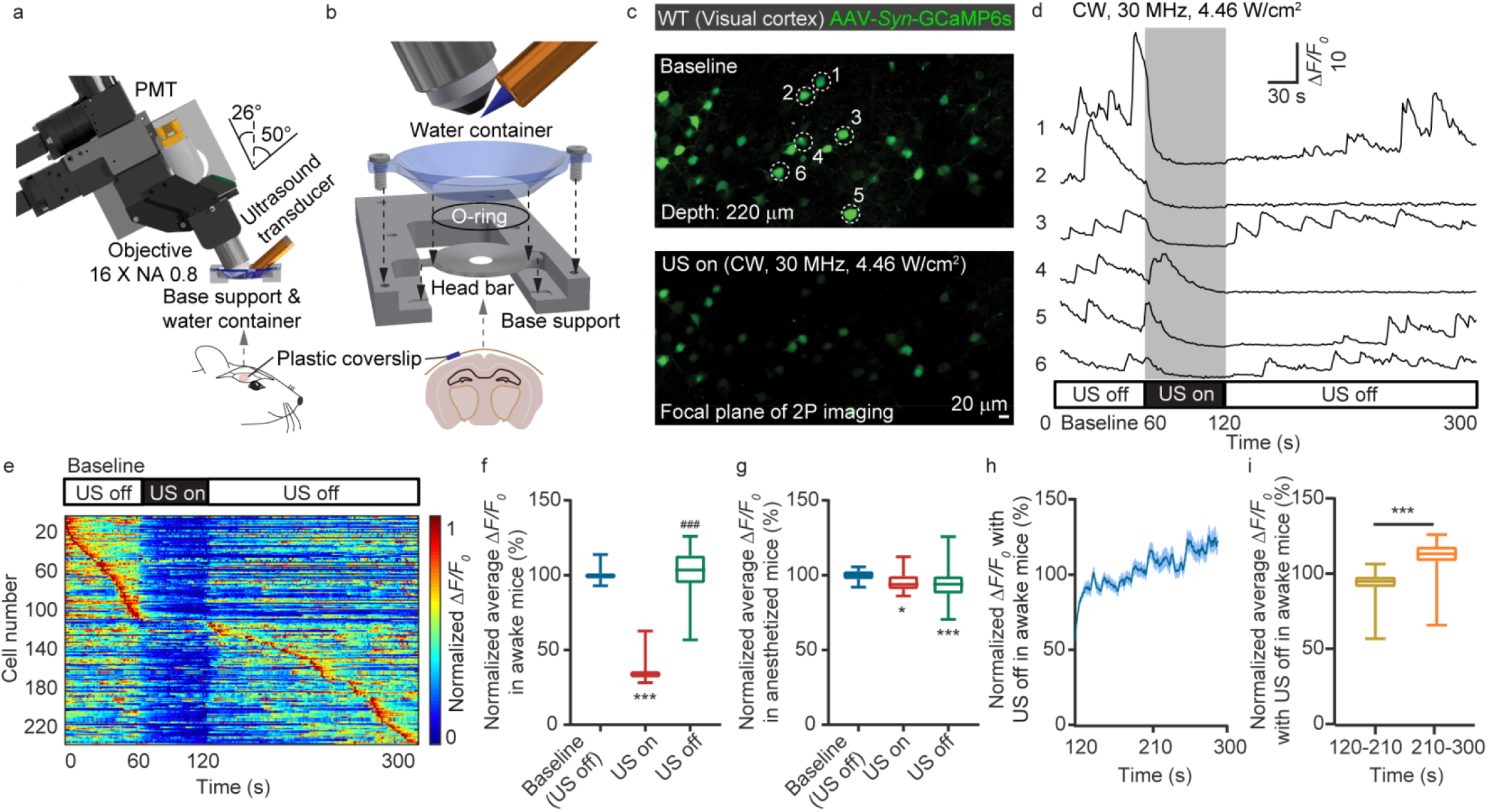
System and implementation of simultaneous ultrasound modulation and two-photon calcium imaging. (a) System design of the dual-modality system combining a two-photon laser scanning fluorescence microscope and a 30 MHz polymer ultrasound transducer. (b) The mechanical assembly including the head bar, O-ring, water container, and base support. (c) Representative two-photon calcium imaging of mouse visual cortex without and with ultrasound modulation. (d) Calcium transient traces for the neurons labeled in **c**. (e) Maximum-normalized calcium activity. (f) Statistics of Δ*F/F*_*0*_ before, during, and after ultrasound modulation for awake mice. (g) Statistics of Δ*F/F*_*0*_ before, during, and after ultrasound modulation for mice in anesthesia. (h) The recovering of the calcium activity in awake mice over time after the ultrasound was turned off. (i) Statistics of Δ*F/F*_*0*_ during the initial and the subsequent 90 seconds after the ultrasound was turned off in awake mice. ** *P* < 0.01, *** or ^###^ *P* < 0.001.

First, we employed the dual-modality system to observe the neuronal response to the 30 MHz ultrasound in the visual cortex of awake mice expressing GCaMP6s. With the ultrasound applied, the calcium transient was significantly reduced (**Fig. 1c-1e**). After the ultrasound was turned off, the calcium transient gradually recovered. To quantify this phenomenon, we performed statistical analysis on the Δ*F/F*_*0*_. The statistics of 4 mice and 180 neurons show that the presence of the 30 MHz ultrasound can reliably suppress the neuronal calcium activity (**Fig. 1f**). However, if the mice were under anesthesia (inherently low neuronal activity and calcium transient), the effect of ultrasound became insignificant (**Fig. 1g**). Although the onset of the ultrasound-induced neural suppression was almost instantaneous, the full recovery of the neuronal activity may take over 100 seconds (**Fig. 1h, 1i**).

Next, we investigated using different ultrasound pressure for calcium transient’s suppression. Experimentally, we repeated the measurements using different signal amplitude to drive the ultrasound transducer and quantified the ultrasound pressure at the sound focus using a calibrated hydrophone (NH0200, Precision Acoustics). For the recorded calcium signals (**Fig. 2a**), we statistically quantified the calcium transients’ frequency, peak Δ*F/F*_*0*_, duration, and integrated transient activity for different sound intensities (**Fig. 2b-2e**). The data show that the calcium transient suppression became visible with 20 V driving signal (2.17 W/cm^2^ at sound focus). With the ultrasound turned off, the calcium activities recovered.

**Figure 2.**
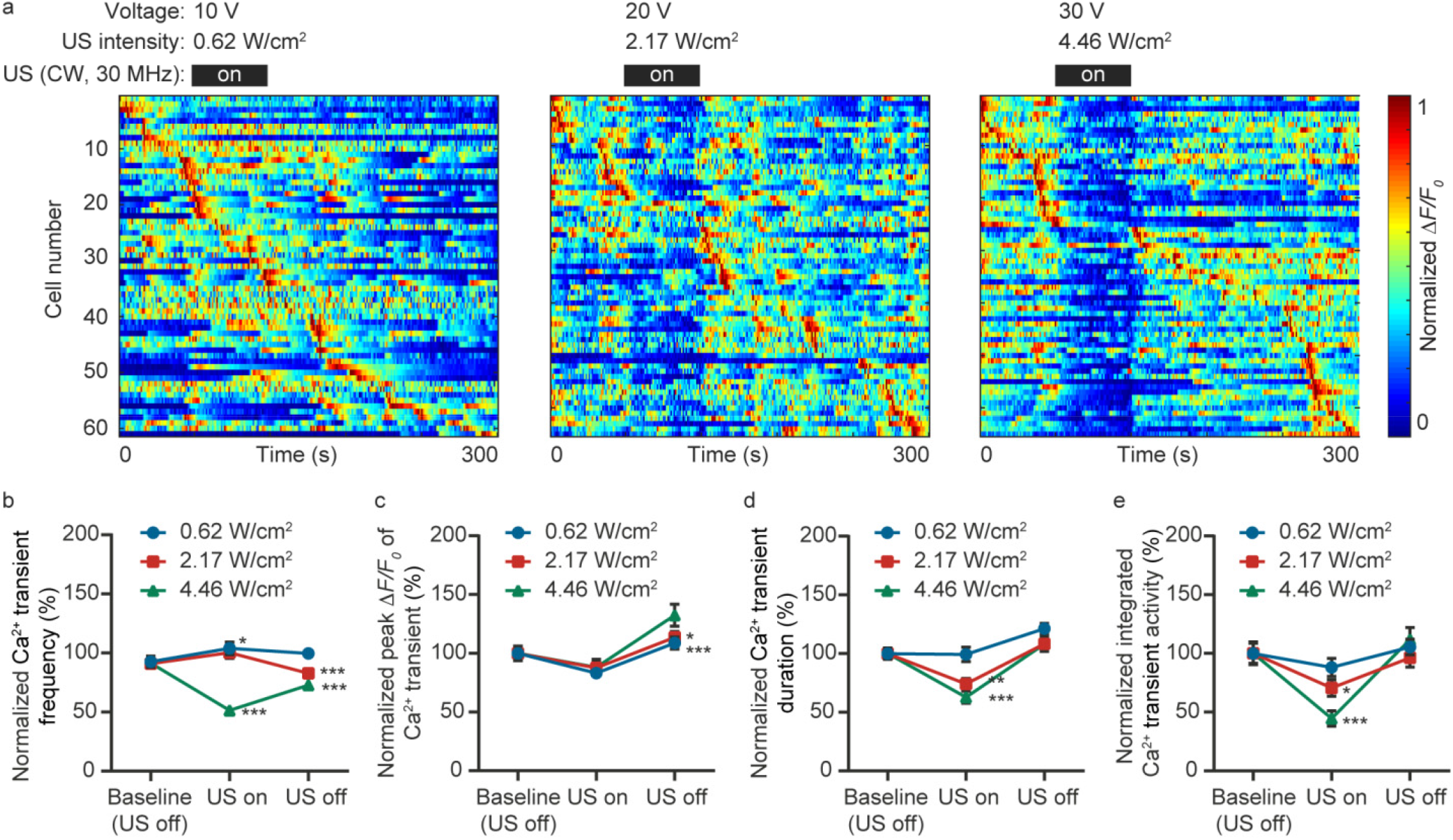
Quantify the ultrasound intensity required for the calcium transient suppression. (a) Maximum-normalized calcium activity measured with different ultrasound intensity. The corresponding applied voltage and ultrasound intensity are shown for each measurement. (b-e) Statistics of the calcium transient frequency, peak Δ*F/F*_*0*_, duration, and integrated activity, respectively. * *P* < 0.05, ** *P* < 0.01, *** *P* < 0.001.

The key advantage of the high-frequency ultrasound is the greatly improved spatial resolution and confinement. The employed polymer transducer featured an element size of 0.25 inch in diameter and a focal length of 0.5 inch, resulting in a focal spot of ∼120 μm in diameter. By spatially translating the brain, we could study the effect of ultrasound on the same neurons and map the ultrasound neural modulation zone (**Fig. 3a, 3b**). Experimentally, a 150 μm horizontal shift could result in an apparent reduction of neuronal modulation effect (**Fig. 3c**), which was comparable to the ultrasound focus spot size.

**Figure 3.**
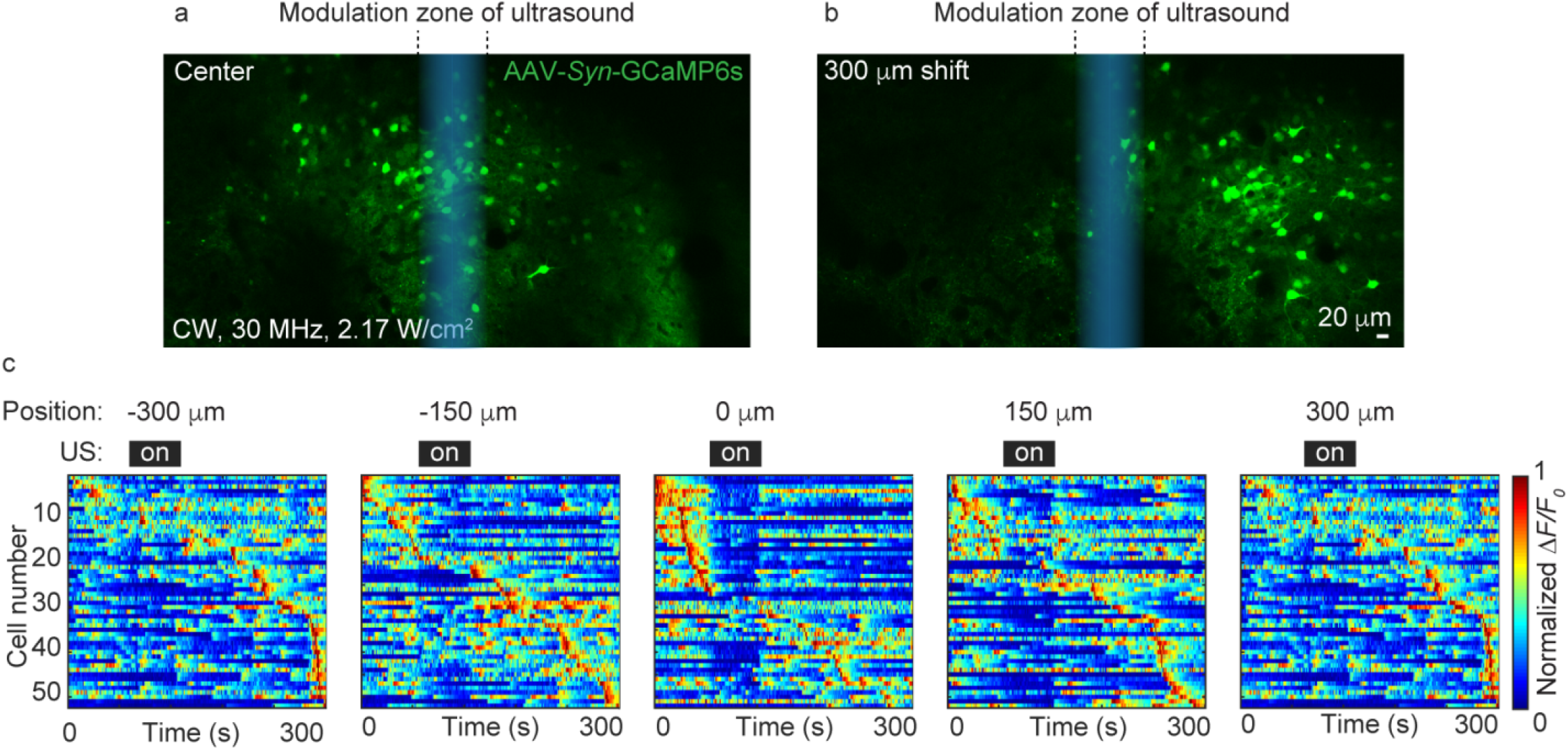
Quantify the spatial resolution of the high-frequency ultrasound neural modulation. (a, b) Representative calcium images before and after the horizontal translation of the brain. (c) Maximum-normalized calcium activity of the same neuronal population measured with different horizontal translation.

Next, we explored the implementation of different carrier frequencies and waveforms. Within the ∼20 MHz bandwidth of our 30 MHz transducer, we tested 20 and 40 MHz ultrasound (**Supplementary Fig. 2a**). Both showed similar suppression effects on neurons albeit the effect of the 40 MHz wave was weaker due to the greater attenuation coefficient in water and the 0.5-inch-long water path. We further tested the effect of amplitude modulation on the 30 MHz ultrasound. Experimentally, we applied pulsed modulation to the 30 MHz carrier signal and varied the duty cycle of the modulation. The data suggest that the reduced duty cycle gradually diminished the effect of neural modulation (**Supplementary Fig. 2b**) and the continuous wave (CW) offered the strongest suppression to calcium transients. For the 10% duty cycle, we also varied the modulation frequency from 100 Hz to 100 kHz, which made a minor difference to the results (**Supplementary Fig. 2c**). Finally, we tested using sinusoidal amplitude modulation and varied the modulation frequency from 0.5 MHz to 2 MHz, which also made little difference. Overall, none of these modulated waveforms worked better than the CW high-frequency ultrasound wave.

As a control experiment, we performed the same modulation measurements on mice that expressed *Thy1*-YFP in neurons. The presence of the 30 MHz ultrasound did not affect the YFP fluorescence signals (**Fig. 4a**). We also tested moving the ultrasound transducer 5 mm away from the imaging FOV. As expected, the effect of the calcium transient suppression disappeared (**Fig. 4b**). The employed ultrasound intensity (2.17-4.46 W/cm^2^) in this study was a bit higher than the common medical ultrasound imaging intensity. To test whether the applied ultrasound could cause any damage to the brain tissue^18^, we applied the 30 MHz ultrasound to the brain of *Cx3cr1*-EGFP mice that expressed EGFP in the microglia. Tissue damage would cause the aggregation of the microglia cells. Although much stronger ultrasound (focus intensity reached 11.8 W/cm^2^) was employed in this control study, no microglia aggregation was observed (**Fig. 4c**).

**Figure 4.**
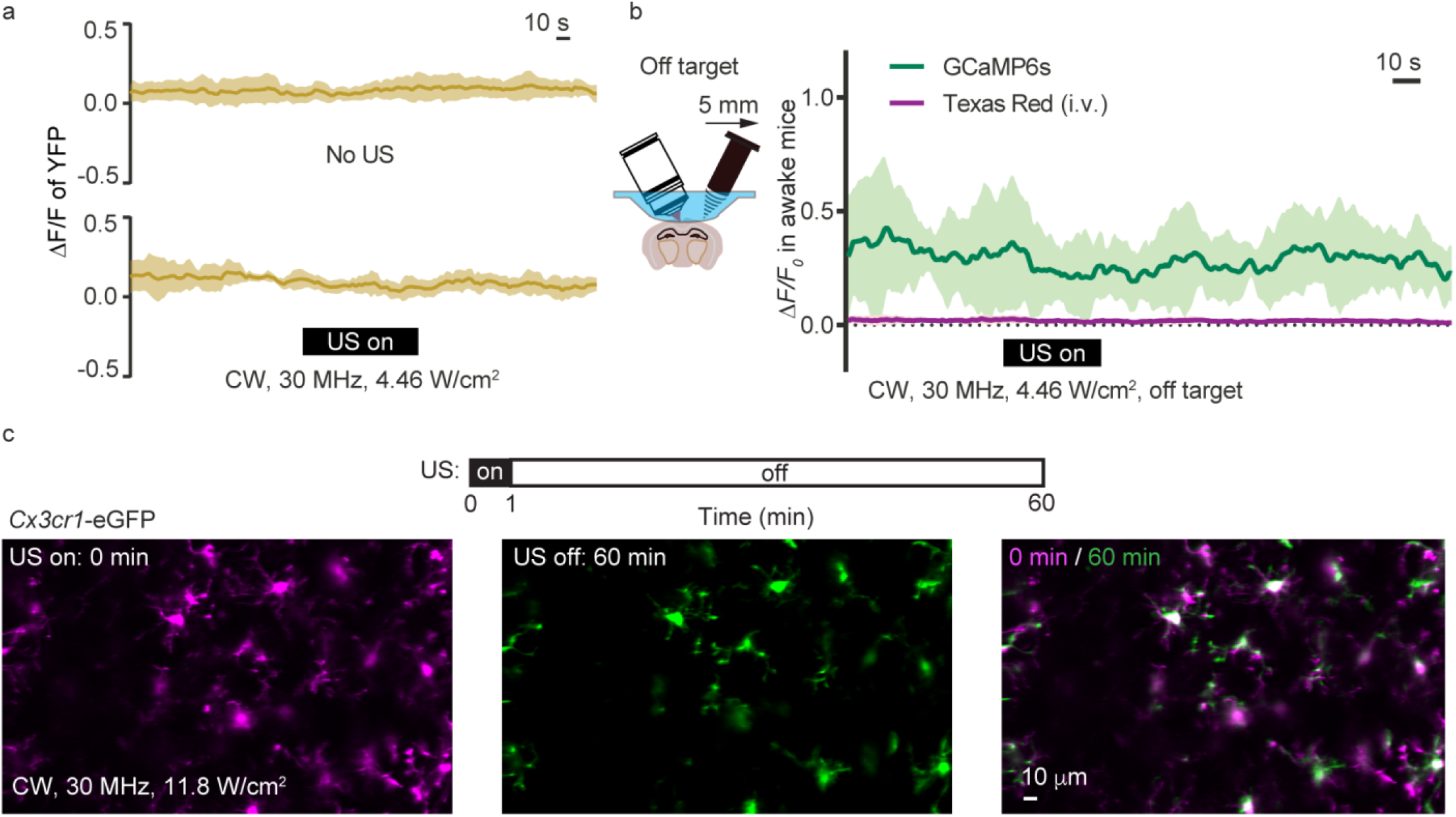
Control studies with YFP, GCaMP6s and Cx3cr1 mice. (a) Fluorescence signal of the *Thy1*-YFP expressing neurons without and with the 30 MHz ultrasound modulation. (b) Calcium imaging with the ultrasound transducer moved 5 mm away from the imaging region. The blood vessels were also visualized by the Texas Red labeling. (c) *Cx3cr1*-EGFP mice imaging with ultrasound modulation. The applied ultrasound intensity (11.8 W/cm^2^) was significantly greater than the typical value used in this study.

In addition to calcium imaging, we further employed Electroencephalogram (EEG) recording to monitor the neuronal population’s response to the high-frequency ultrasound (**Fig. 5a**). Using wavelet transform, we decomposed the EEG signals into different frequency ranges (**Fig. 5b**). We found that the low frequency (0.3-1.3 Hz) activity was enhanced by the ultrasound wave while the high-frequency components largely remained the same (**Fig. 5c**). As a control measurement, we moved the ultrasound transducer 5 mm away from the EEG recording site. As expected, the EEG signal variation vanished (**Fig. 5d**). Next, we studied the EEG signal variation as a function of the applied ultrasound intensity. The data suggest that noticeable signal variation appeared when the ultrasound intensity rose above ∼2 W/cm^2^ (**Fig. 5e**), which agreed well with the two-photon calcium imaging results (**Fig. 2a**). Interestingly, the EEG signal variation would again disappear if the mice were under anesthesia (**Fig. 2f**), very similar to the observation by the calcium imaging (**Fig. 1g**).

**Figure 5.**
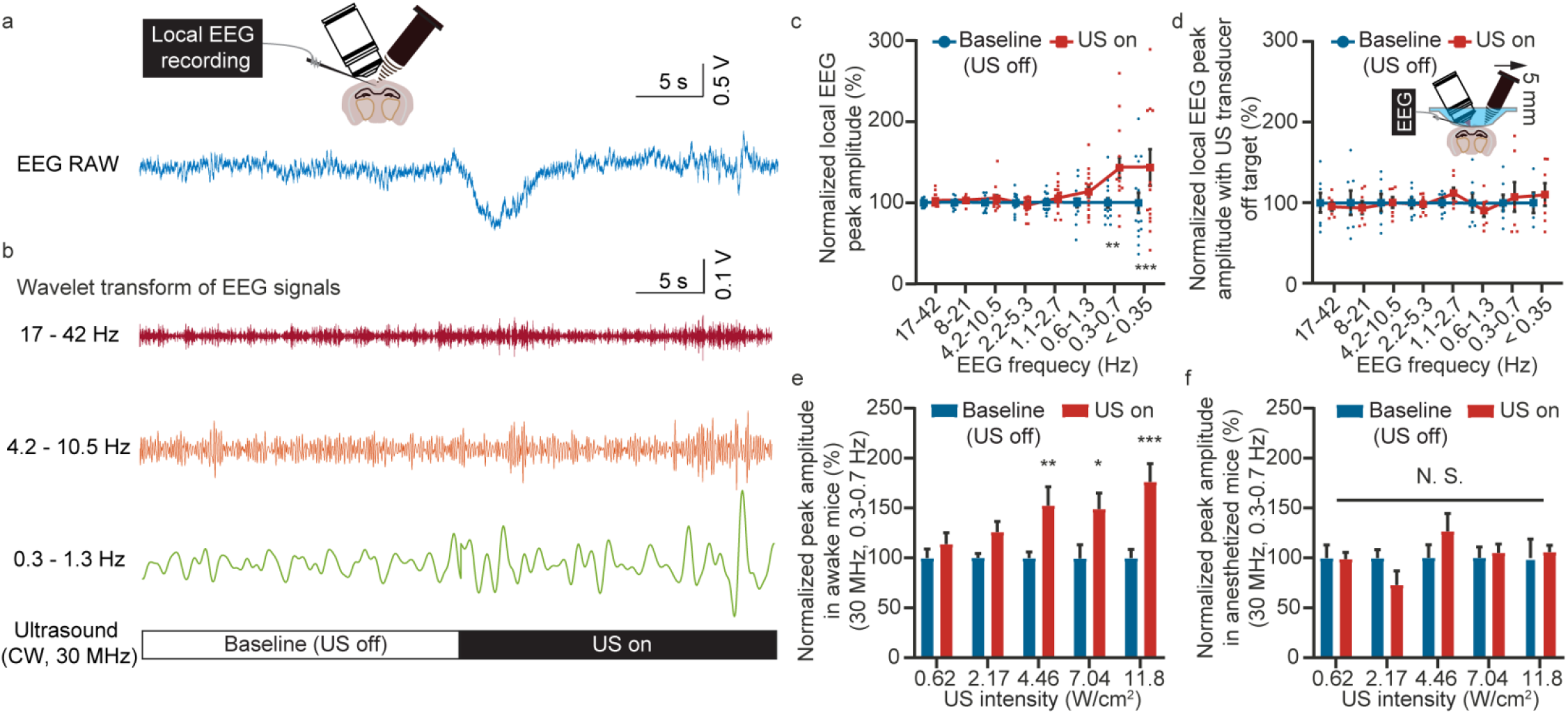
Simultaneous EEG recording during high-frequency ultrasound modulation. (a) Representative raw data of the EEG recording. (b) EEG signals decomposed to different frequency range by wavelet transform. (c) Statistics of the EEG peak amplitude as a function of frequency range before and after the ultrasound modulation. (d) Statistics of the EEG peak amplitude as a function of frequency before and after the ultrasound modulation with the ultrasound transducer moved 5 mm away. (e) Statistics of the peak amplitude for 0.3-0.7 Hz EEG signal with ultrasound of different intensity applied to awake mice. (f) Statistics of the peak amplitude for 0.3-0.7 Hz EEG signal with ultrasound of different intensity applied to anesthetized mice. * *P* < 0.05, ** *P* < 0.01, *** *P* < 0.001.

## Discussion

The data from both the calcium imaging and the EEG recording suggest that the high frequency (∼30 MHz) ultrasound can modulate neuronal activities, effectively shrinking the neural modulation volume by a factor of 27,000. This will provide significantly improved resolutions and degrees of freedom for neural modulation. The spatial confinement of the ultrasound modulation in this work was limited by the numerical aperture (NA = 0.24) of the off-the-shelf transducer. Further improvement on the focusing lenses (e.g. customizing NA 0.5 ultrasound lenses) may potentially achieve ∼50 μm and ∼300 μm in the transverse and axial resolutions, respectively. Such resolutions would be highly valuable for modulating individual function regions in the small rodents’ brains.

Limited by the transducer bandwidth, the highest ultrasound frequency applied was 40 MHz^39^. To achieve even greater spatial resolution, higher frequencies (e.g. 60 MHz) should be tested in future studies, which could further reduce the modulation volume by another factor of 8. Although the water attenuation will lead to a shorter working distance (e.g. 5-6 mm) in brain tissue, it would still be sufficient to cover the majority of the mouse brain volume.

Compared to the common 1 MHz ultrasound^40,41^, a key limitation of the 30 MHz ultrasound is that it will suffer from stronger loss through thick skulls (e.g. human skull)^5,42–44^. Therefore, it may not be able to achieve noninvasive transcranial performance on the human brain^45–47^. However, the attenuation through the mouse skull (**Supplementary Table 1**) is moderate and can be compensated by increasing the transducer input power. Therefore, high-frequency neural modulation can be implemented on mouse brain noninvasively. For bigger mammal models with thick skulls, the alternative solution is to perform surgery to replace part of the skull with plastic that matches the brain tissue in acoustic impedance, similar to the plastic cranial window with ∼90% power transmission employed in this work.

A constraint in the employed dual-modality system is that the ultrasound focus was only applied to one location. Changing the ultrasound focus required manually adjusting a 3-axis linear stage. Potentially, customized high-frequency ultrasound transducer arrays should be employed to achieve arbitrary simultaneous multi-foci 3D neural modulation.

In summary, we present the experimental results acquired by a dual-modality system combining two-photon calcium imaging and high-frequency focused ultrasound. The calcium imaging data suggest that the 30 MHz ultrasound can effectively suppress the calcium transients in awake mice. However, such an effect would disappear when the mice were under anesthesia. Additionally, the EEG measurement showed that the ultrasound increased the low-frequency EEG signals. The effect would disappear when the mice were under anesthesia, similar to the observation by the two-photon calcium imaging. Moreover, the minimum ultrasound intensity (∼2 W/cm^2^) required to cause EEG signal variation also agreed well with that of the calcium measurement. As a variety of neurological disorders such as epilepsy^48,49^, Alzheimer’s disease^50–52^, and neuropathic pain^53–56^ are associated with hyperactive neuronal activities, the capabilities of noninvasively suppressing neuronal responses are highly desirable for treatment and symptom alleviation. The *in vivo* results revealed in this study indicate that the ∼30 MHz ultrasound can be employed for reliable neural suppression, paving the way for high-resolution neural activity control in neuroscience research and potential medical applications. Moreover, the dual-modality system developed and validated in this study can serve as a general platform for studying neuronal, glial, and vasculature dynamics induced by various ultrasound modulations.

## Methods

### Animal preparation

The wild type C57BL/6 mice, the *Thy1*-YFP mice, and the *Cx3cr1*-EGFP mice were all purchased from the Jackson Laboratories. For calcium imaging, we injected AAV1-*Syn*-GCaMP6s-WPRE375-SV40 virus (Addgene, 100843-AAV1, 1×10^13^) into the visual cortex of the C57BL/6 mice 2-3 weeks before the *in vivo* imaging experiments. The cranial window surgeries were performed a few days before the imaging. The exposed cortex was covered by a 3 mm diameter plastic coverslip that was attached to the skull. For the vascular labeling, Dextran & Texas Red (70,000 MW, 5%, 50 mg/kg, Thermo Fisher) was employed through orbital injection. All procedures involving mice were approved by the Animal Care and Use Committees of Purdue University.

### Installation of the water container

Different from the common calcium imaging implementation, we need to support a long water path for the 30 MHz transducer which featured a 0.5-inch-long working distance. This involves custom machined base support, large diameter head bar (1.8 g in weight), O-ring (0.5 mm and 32 mm in height and diameter, respectively), and water container (**Supplementary Fig. 3**). First, we put the extrusions of the head bar into the indentation on the base support. Next, we placed the O-ring on the outer edge of the head bar, then pressed the O-ring onto the head bar with the water container above, and finally locked the container position using two low-profile 8-32 machine screws. The pressed O-ring ensured a watertight connection from the head bar to the water container. The base support was attached to a 3-axis motorized stage (LTA-HL, Newport) which precisely positioned the mouse under the imaging system.

### *In vivo* two-photon imaging and data analysis

The calcium imaging data were primarily collected from layer 2/3 neurons of the visual cortex. We also tested the motor cortex which showed the same effects (**Supplementary Fig. 4**). The calcium image recording rate was 4 Hz. The typical recording session was 300 sec long. In the first 60 sec, the ultrasound was off (recording baseline activity). In the next 60 sec, the ultrasound was on (modulation). In the last 180 sec, the ultrasound was off (recovery). Before extracting the calcium transients from the somata, we utilized the average of the time-lapsed images as the position reference and performed spatial cross-correlations for motion correction. We employed normalized heatmaps to assist the visualization of neuronal activity (e.g. **Fig. 1e, 2a, 3c**), in which the calcium activity (Δ*F/F*_*0*_) of each neuron was normalized by its maximum value and the neurons were ordered according to the time of maximum Δ*F/F*_*0*_ responses. The minimum 10% of each cell’s fluorescence signal was defined as *F*_*0*_. The threshold for detecting calcium transient was for Δ*F/F*_*0*_ > 0.5. The full width at half maximum (FWHM) of each calcium transient was measured as the calcium transient duration^57^. The frequency was defined as the number of calcium transients per second and the peak value was defined as the maximum value within each calcium transient. The total calcium activity was defined as the accumulated area under the curve per second.

### Electroencephalogram (EEG) recording and data analysis

EEG signals were recorded by using an integrated data acquisition and analysis system (BL-420F, TME technology). The electrode was made from an epoxy coated silver wire (0.005 inches in diameter). The electrodes were carefully inserted into the V1 cortex which was then covered by a plastic coverslip. Before the EEG recording, we translated the brain under the two-photon microscope such that the electrodes were at the center of the focal plane to ensure that the electrodes were near the ultrasound modulation zone. The wavelet-based signal decomposition was implemented by using the wavelet toolbox of MATLAB (MathWorks).

### Statistics

All the data in this study are represented as mean ± standard error. All the data analysis was performed with the Shapiro-Wilk normality test. For the data that passed the normality test (*P* > 0.05), an unpaired t-test with Welch’s correction was selected for two groups’ comparison, the one or two-way ANOVA Tukey’s or Sidak’s multiple comparisons test were selected for comparing multiple groups. *P* < 0.05 is recognized as statistically significant. All statistical analyses were performed using *GraphPad Prism*. No results of the successful acquisition from images and measurements were excluded and filtered. The experiment did not include randomized and blinding experiments. At least three mice were used in each experiment. *P* values, *n*, and the statistical tests for all the experiments were summarized in **supplementary table 2**.

## Acknowledgment

This work was funded by NIH (U01NS118302, U01NS107689) and Purdue University. M.C. thanks Howard Hughes Medical Institute for scientific instruments. Q.Z. acknowledges the support by NIH R01EY030126.

## Author Contributions

M.C. conceived the study, designed, and implemented the dual-modality system. M.C., W.G., and Q.Z. supervised the research. Z.C. designed and carried out the biology experiment and performed data analysis and statistics. C.W. quantified the ultrasound intensity and pressure. B.W. performed the wavelet analysis for the EEG recording. Z.C. C.W. and B.W. collaborated on the figure preparation. All authors contributed to the writing of the manuscript.

## Data availability

The datasets generated during and/or analyzed during the current study are available from the corresponding author on reasonable request.

## Correspondence

Correspondence and requests for materials should be addressed to M.C.

