## Supplementary Material for "High resolution ultrasonic neural modulation observed via *in vivo* two-photon calcium imaging"

**This file includes Supplementary Figure 1-4 and Supplementary Table 1-2.**

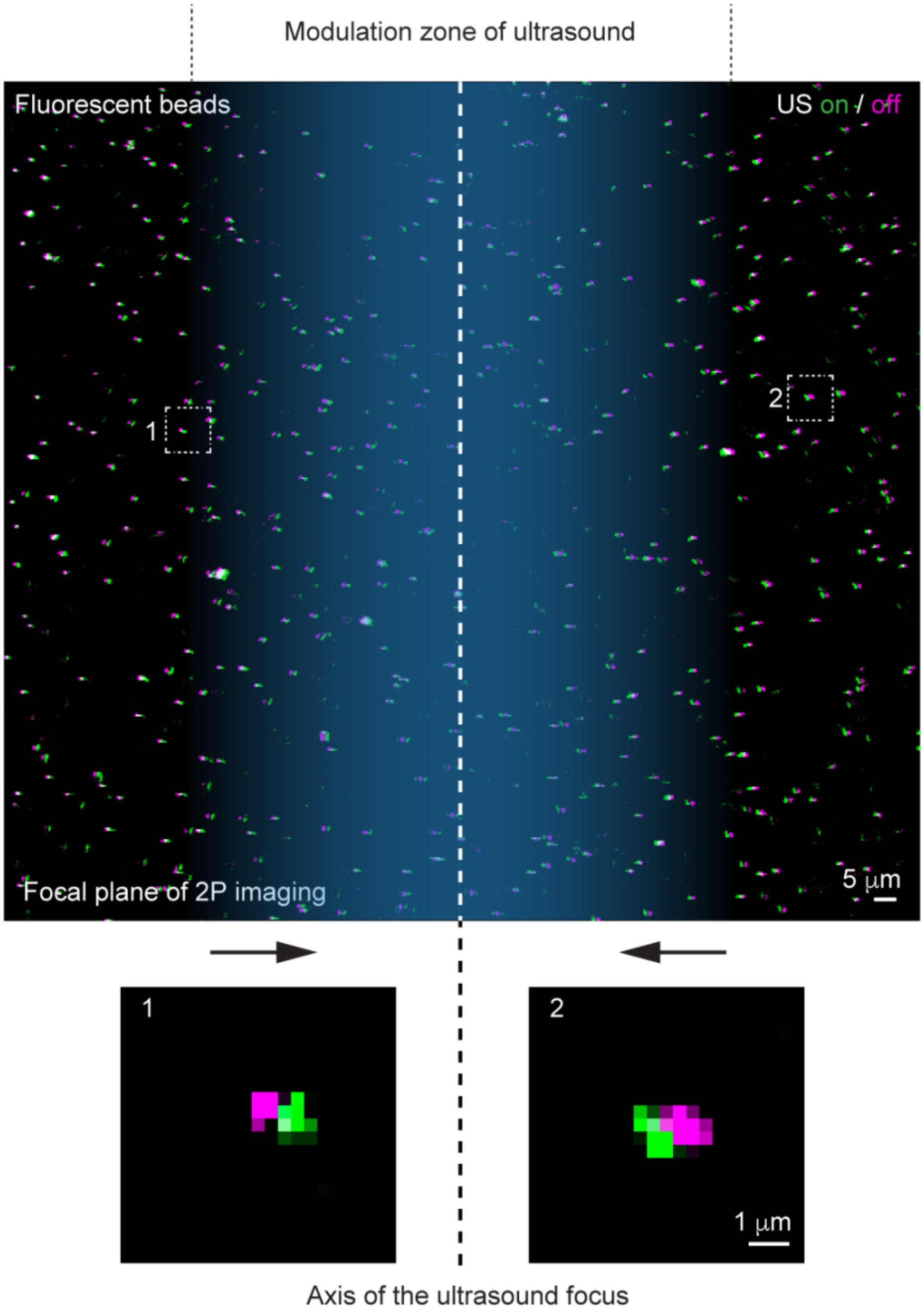


**Supplementary Figure 1 | Procedure for visualizing the ultrasound focus via the two-photon laser scanning fluorescence microscope**. The propagation of the ultrasound resulted in a low-pressure zone near the focus, which pulled the fluorescence beads in agar towards the ultrasound focus by hundreds of nanometers. Showing the two images recorded with the ultrasound on and off in two pseudo colors allowed us to visualize the ultrasound focus location.

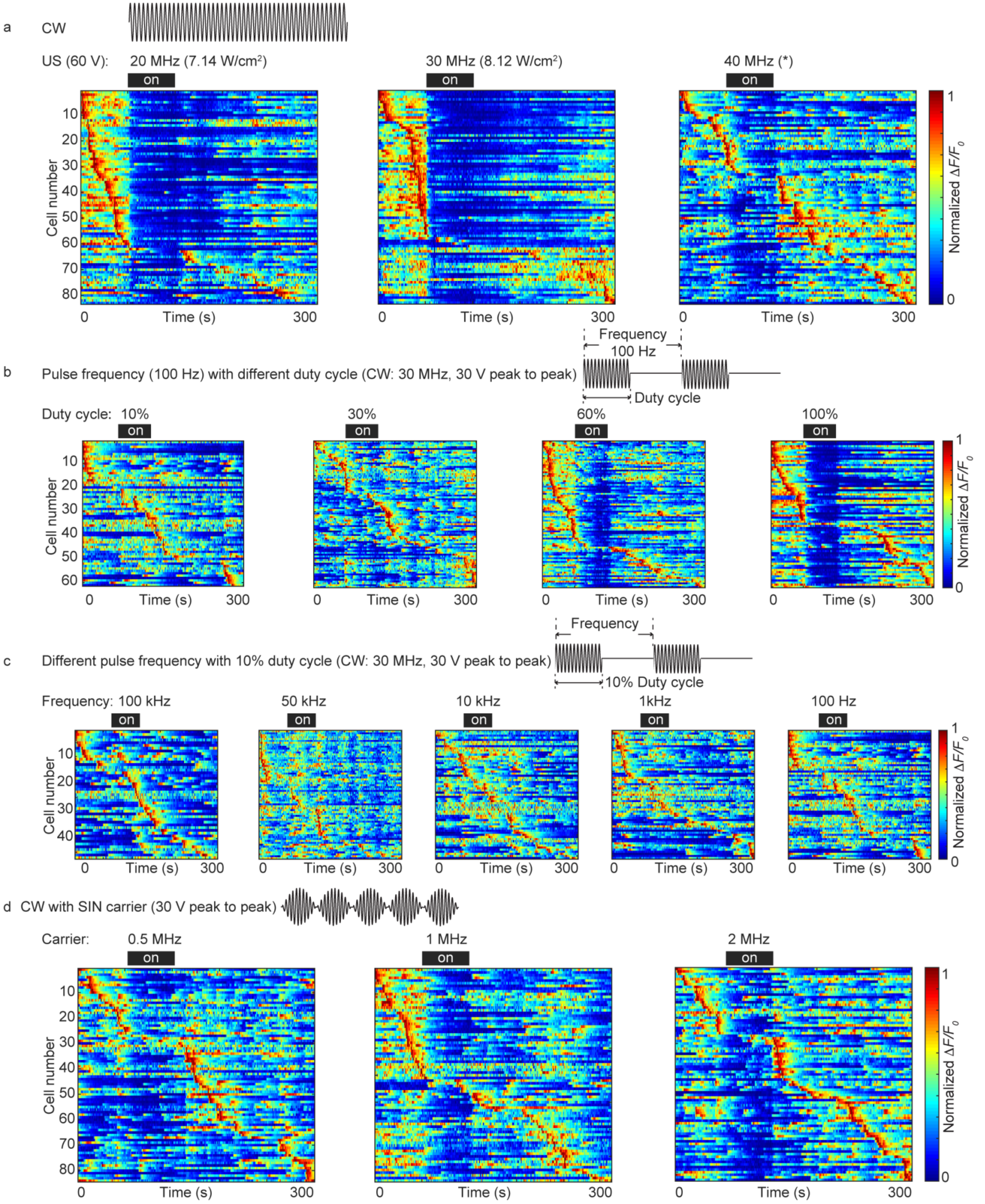

a
CW
US (60 V):
20 MHz (7.14 W/cm²)
30 MHz (8.12 W/cm²)
40 MHz (*)
on
Cell number
Time (s)
0
300
Normalized ΔF/F0
b
Pulse frequency (100 Hz) with different duty cycle (CW: 30 MHz, 30 V peak to peak)
Frequency
100 Hz
Duty cycle
Duty cycle: 10%
30%
60%
100%
c
Different pulse frequency with 10% duty cycle (CW: 30 MHz, 30 V peak to peak)
Frequency
10% Duty cycle
Frequency: 100 kHz
50 kHz
10 kHz
1kHz
100 Hz
d
CW with SIN carrier (30 V peak to peak)
Carrier:
0.5 MHz
1 MHz
2 MHz

**Supplementary Figure 2 | Dependence of calcium transient suppression on the ultrasound carrier frequency, pulsed modulation duty cycle, pulsed modulation frequency, and sinusoidal modulation frequency**. (a) Maximum-normalized calcium activity for the same neuronal population with 20, 30, and 40 MHz ultrasound applied in the measurement. The applied driving signal was 60 v in all three data sets. The measured (by calibrated hydrophone) ultrasound intensity values are shown. The 40 MHz signal was outside the calibration range of the hydrophone. (b) Calcium activity with the pulsed amplitude modulation of different duty cycles. The CW wave showed the strongest effect on suppressing calcium transients. (c) Calcium activity with different pulsed amplitude modulation frequencies (from 100 Hz to 100 kHz). The duty cycle was 10%. (d) Calcium activity with different sinusoidal amplitude modulation frequencies (from 0.5 MHz to 2 MHz).

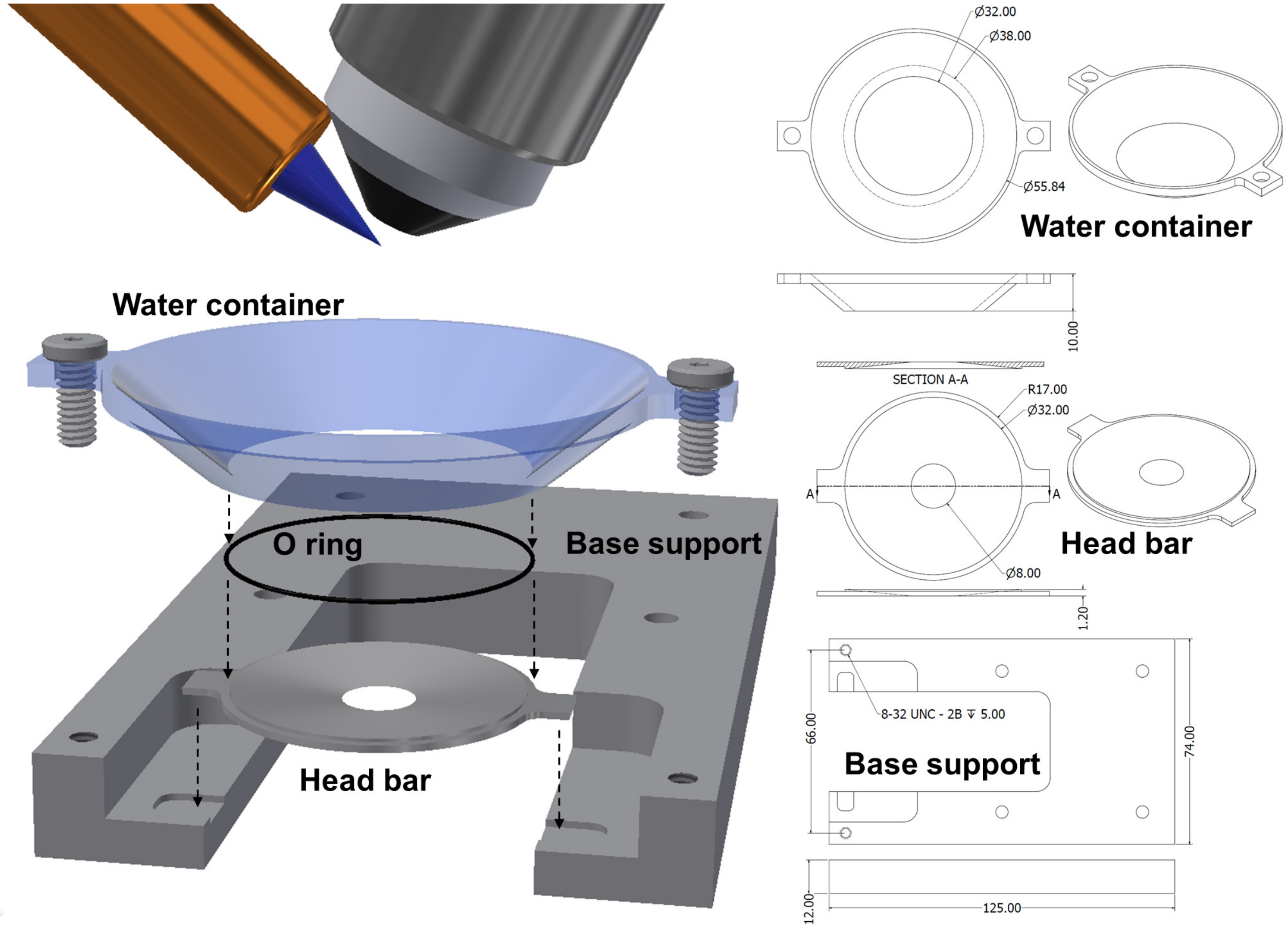


**Supplementary Figure 3 | Mechanical design of the watertight mouse head bar support system**. The dimensions of the water container, head bar, and base support are shown. The O ring was 0.5 mm and 32 mm in thickness and diameter, respectively. The dimension unit is mm. All parts were fabricated with 6061 aluminum alloy. The head bar weight was 1.8 g.

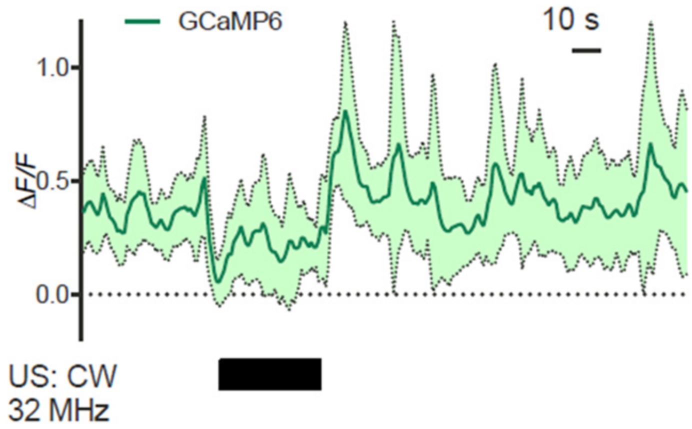


**Supplementary Figure 4 | Ultrasound modulation to the motor cortex of awake mice.** The calcium image recording employed was the same as that of Fig.1. The difference was that the measurement location was the motor cortex. The calcium transients and the statistics were from three mice.

**Supplementary Table 1 | Calibrated hydrophone measured ultrasound focal pressure and intensity**.

| | | 20 V | | 30 V | | 40 V | |
|---|---|---|---|---|---|---|---|
| | | MPa | W/cm$^2$ | MPa | W/cm$^2$ | MPa | W/cm$^2$ |
| 20MHz | None | 0.50 | 2.11 | 0.74 | 4.57 | 0.97 | 7.97 |
| | Plastic | 0.46 | 1.76 | 0.69 | 4.04 | 0.91 | 7.03 |
| | Skull | 0.31 | 0.81 | 0.51 | 2.24 | 0.69 | 4.04 |
| 30MHz | None | 0.53 | 2.32 | 0.75 | 4.67 | 1.02 | 8.69 |
| | Plastic | 0.51 | 2.17 | 0.73 | 4.46 | 0.94 | 7.40 |
| | Skull | 0.36 | 1.06 | 0.53 | 2.32 | 0.70 | 4.11 |

We show the measured values in water only, through the plastic cranial window, and through a piece of mouse skull. In the measurement, the plastic cranial window and the skull were held along the horizontal plane as the case of the actual imaging configuration.

**Supplementary Table 2 | Statistics of experimental data.**

| Fig. | Items | Mean | s.e.m | *n* | Mice | Statistical test | *P* value |
|---|---|---|---|---|---|---|---|
| 1f | Baseline | 100.00 | 0.22 | 180 | 4 | Dunnett's multiple comparisons test | - |
| | US on | 35.07 | 0.36 | 180 | | | <0.0001 |
| | US off | 103.30 | 0.46 | 180 | | | 0.0009 |
| 1g | Baseline | 100 | 0.4868 | 217 | 4 | Dunnett's multiple comparisons test | - |
| | US on | 95.56 | 0.9459 | 217 | | | 0.0106 |
| | US off | 94.2 | 0.7584 | 217 | | | <0.0001 |
| 1i | 120-210 | 93.92 | 0.4257 | 180 | 4 | Unpaired t test | - |
| | 210-300 | 112.7 | 0.4159 | 180 | | | <0.0001 |
| 2b | Baseline 0.62 | 100.00 | 4.44 | 121 | 4 | Tukey's multiple comparisons test | - |
| | US on 0.62 | 110.85 | 5.17 | 126 | | | 0.0465 |
| | US off 0.62 | 84.91 | 3.32 | 156 | | | 0.0733 |
| | Baseline 2.17 | 100.00 | 3.71 | 178 | | | - |
| | US on 2.17 | 107.34 | 4.70 | 184 | | | 0.1391 |
| | US off 2.17 | 80.98 | 2.84 | 199 | | | 0.0001 |
| | Baseline 4.46 | 100.00 | 3.62 | 148 | | | - |
| | US on 4.46 | 71.28 | 3.88 | 111 | | | <0.0001 |
| | US off 4.46 | 76.28 | 3.86 | 152 | | | <0.0001 |
| 2c | Baseline 0.62 | 100.00 | 6.27 | 121 | 4 | Tukey's multiple comparisons test | - |
| | US on 0.62 | 83.07 | 3.64 | 126 | | | 0.0663 |
| | US off 0.62 | 108.81 | 5.18 | 156 | | | 0.5606 |
| | Baseline 2.17 | 100.00 | 5.81 | 178 | | | - |
| | US on 2.17 | 87.52 | 5.42 | 184 | | | 0.1697 |
| | US off 2.17 | 113.51 | 5.14 | 199 | | | 0.0492 |
| | Baseline 4.46 | 100.00 | 5.62 | 148 | | | - |
| | US on 4.46 | 88.07 | 6.77 | 111 | | | 0.718 |
| | US off 4.46 | 132.49 | 9.41 | 152 | | | <0.0001 |
| 2d | Baseline 0.62 | 100.00 | 4.89 | 121 | 4 | Tukey's multiple comparisons test | - |
| | US on 0.62 | 99.26 | 6.21 | 126 | | | 0.9946 |
| | US off 0.62 | 121.22 | 4.49 | 156 | | | 0.0001 |
| | Baseline 2.17 | 100.00 | 4.84 | 178 | | | - |
| | US on 2.17 | 74.02 | 4.66 | 184 | | | <0.0001 |
| | US off 2.17 | 108.29 | 4.96 | 199 | | | 0.3176 |
| | Baseline 4.46 | 100.00 | 4.76 | 148 | | | - |
| | US on 4.46 | 62.73 | 5.16 | 111 | | | <0.0001 |
| | US off 4.46 | 108.15 | 6.30 | 152 | | | 0.5049 |
| 2e | Baseline 0.62 | 100.00 | 8.83 | 121 | 4 | Tukey's multiple comparisons test | - |
| | US on 0.62 | 88.14 | 7.80 | 126 | | | 0.6022 |
| | US off 0.62 | 105.47 | 6.80 | 156 | | | 0.8813 |
| | Baseline 2.17 | 100.00 | 9.80 | 178 | | | - |
| | US on 2.17 | 70.54 | 7.03 | 184 | | | 0.0266 |
| | US off 2.17 | 96.23 | 7.91 | 199 | | | 0.9799 |
| | Baseline 4.46 | 100.00 | 8.54 | 148 | | | - |
| | US on 4.46 | 44.56 | 6.48 | 111 | | | <0.0001 |
| | US off 4.46 | 111.73 | 10.38 | 152 | | | 0.6799 |
| 5c | Baseline 17-42 | 100.00 | 1.16 | 15 | 3 | Sidak's multiple comparisons test | - |
| | Baseline 8-21 | 100.00 | 1.55 | 15 | | | - |
| | Baseline 4.2-10.5 | 100.00 | 2.08 | 15 | | | - |
| | Baseline 2.2-5.3 | 100.00 | 2.94 | 15 | | | - |
| | Baseline 1.1-2.7 | 100.00 | 2.84 | 15 | | | - |
| | Baseline 0.6-1.3 | 100.00 | 5.43 | 15 | | | - |

| | | | | | | | |
|---|---|---|---|---|---|---|---|
| | Baseline 0.3-0.7 | 100.00 | 5.86 | 15 | | | - |
| | Baseline < 0.35 | 100.00 | 12.50 | 15 | | | - |
| | US on 17-42 | 102.47 | 2.06 | 15 | | | >0.9999 |
| | US on 8-21 | 102.92 | 1.18 | 15 | | | >0.9999 |
| | US on 4.2-10.5 | 105.43 | 3.83 | 15 | | | 0.9996 |
| | US on 2.2-5.3 | 96.63 | 2.87 | 15 | | | >0.9999 |
| | US on 1.1-2.7 | 105.75 | 4.24 | 15 | | | 0.9994 |
| | US on 0.6-1.3 | 113.57 | 6.69 | 15 | | | 0.8693 |
| | US on 0.3-0.7 | 143.27 | 12.50 | 15 | | | 0.0011 |
| | US on < 0.35 | 143.74 | 22.37 | 15 | | | 0.0009 |
| 5d | Baseline 17-42 | 100.00 | 12.18 | 7 | 3 | Sidak's multiple comparisons test | - |
| | Baseline 8-21 | 100.00 | 15.13 | 7 | | | - |
| | Baseline 4.2-10.5 | 100.00 | 10.78 | 7 | | | - |
| | Baseline 2.2-5.3 | 100.00 | 6.94 | 7 | | | - |
| | Baseline 1.1-2.7 | 100.00 | 5.50 | 7 | | | - |
| | Baseline 0.6-1.3 | 100.00 | 11.96 | 7 | | | - |
| | Baseline 0.3-0.7 | 100.00 | 9.89 | 7 | | | - |
| | Baseline < 0.35 | 100.00 | 12.56 | 5 | | | - |
| | US on 17-42 | 95.46 | 5.08 | 7 | | | >0.9999 |
| | US on 8-21 | 93.94 | 7.90 | 7 | | | >0.9999 |
| | US on 4.2-10.5 | 100.29 | 6.74 | 7 | | | >0.9999 |
| | US on 2.2-5.3 | 98.24 | 4.82 | 7 | | | >0.9999 |
| | US on 1.1-2.7 | 111.83 | 6.90 | 7 | | | 0.8424 |
| | US on 0.6-1.3 | 90.37 | 7.64 | 7 | | | 0.9968 |
| | US on 0.3-0.7 | 107.03 | 18.26 | 7 | | | >0.9999 |
| | US on < 0.35 | 110.24 | 14.07 | 7 | | | 0.9993 |
| 5e | Baseline 0.62 | 100.00 | 8.78 | 10 | 3 | Sidak's multiple comparisons test | - |
| | Baseline 2.17 | 100.00 | 4.44 | 13 | | | - |
| | Baseline 4.46 | 100.00 | 5.86 | 15 | | | - |
| | Baseline 7.04 | 100.00 | 13.62 | 6 | | | - |
| | Baseline 11.8 | 100.00 | 8.45 | 9 | | | - |
| | US on 0.62 | 113.77 | 11.55 | 10 | | | 0.9533 |
| | US on 2.17 | 125.90 | 10.73 | 13 | | | 0.0552 |
| | US on 4.46 | 152.53 | 18.88 | 15 | | | 0.0038 |
| | US on 7.04 | 149.14 | 15.73 | 6 | | | 0.0389 |
| | US on 11.8 | 176.36 | 18.16 | 9 | | | 0.0008 |
| 5f | Baseline 0.62 | 100.00 | 13.02 | 4 | 3 | Sidak's multiple comparisons test | - |
| | Baseline 2.17 | 100.00 | 8.33 | 3 | | | - |
| | Baseline 4.46 | 100.00 | 13.17 | 4 | | | - |
| | Baseline 7.04 | 100.00 | 10.91 | 4 | | | - |
| | Baseline 11.8 | 100.00 | 18.77 | 4 | | | - |
| | US on 0.62 | 98.94 | 6.65 | 4 | | | >0.9999 |
| | US on 2.17 | 72.83 | 14.04 | 3 | | | 0.5835 |
| | US on 4.46 | 126.84 | 17.61 | 4 | | | 0.5175 |
| | US on 7.04 | 105.09 | 8.79 | 4 | | | 0.9994 |
| | US on 11.8 | 106.03 | 6.60 | 4 | | | 0.9986 |
